# Untwisted α-synuclein Filaments formed in the Presence of Lipid Vesicles

**DOI:** 10.1101/2021.10.19.464986

**Authors:** Anvesh K. R. Dasari, Lucas Dillard, Alimohammad Hojjatian, Urmi Sengupta, Rakez Kayed, Kenneth A. Taylor, Mario J. Borgnia, Kwang Hun Lim

## Abstract

Accumulation of filamentous aggregates of α-synuclein is a pathological hallmark of several neurodegenerative diseases including Parkinson’s disease (PD). Interaction between α-synuclein and lipids has been shown to play a critical role in aggregation of α-synuclein. Most structural studies have, however, been focused on α-synuclein filaments formed in the absence of lipids. Here, we report structural investigation of α-synuclein filaments assembled under the quiescent conditions in the presence of anionic lipid vesicles using electron microscopy (EM) including cryo-EM. Our transmission electron microscopy (TEM) analyses reveal that α-synuclein forms curly protofilaments at an early stage of aggregation. The flexible protofilaments were then converted to long filaments after a longer incubation of 30 days. More detailed structural analyses using cryo-EM reveal that the long filaments adopt untwisted structures with different diameters, which have not been observed in previous α-synuclein filaments formed in vitro. The untwisted filaments are rather similar to straight filaments with no observable twist that are extracted from patients with dementia with Lewy bodies. Our structural studies highlight the conformational diversity of α-synuclein filaments, requiring additional structural investigation of not only more ex vivo α-synuclein filaments, but also in vitro α-synuclein filaments formed in the presence of diverse co-factors to better understand the molecular basis of diverse molecular conformations of α-synuclein filaments.

## Introduction

Intracellular deposition of misfolded α-synuclein is a hallmark of Parkinson’s disease (PD), dementia with Lewy bodies (DLB), and multiple system atrophy (MSA), collectively termed synucleinopathies.^1^ Structural characterization of filamentous α-synuclein aggregates is essential to understanding the molecular mechanism of α-synuclein aggregation. Atomic-resolution structures of α-synuclein filaments solved by solid-state NMR and cryo-EM revealed diverse molecular conformations,^2–6^ suggesting the presence of multiple misfolding and aggregation pathways. Recent structural studies of ex vivo α-synuclein filaments revealed that α-synuclein filaments extracted from MSA and DLB patients’ brains exhibit distinct molecular conformations from those of the in vitro α-synuclein filaments.^7^ In particular, filaments derived from DLB patient’s brains were shown to adopt untwisted fibril morphologies,^7^ which have not been observed for in vitro α-synuclein filaments^8^. In addition, our recent structural studies showed that interaction between an aggregation-prone protein, tau, and α-synuclein promotes the formation of α-synuclein filaments with distinct conformations.^9,10^ Previous solid-state NMR studies also revealed that α-synuclein filaments derived by anionic phospholipids have distinct molecular conformations in the N-terminal region.^11^ These results suggest that cofactors in cellular environments might interact with α-synuclein, leading the protein to a specific misfolding pathway for the formation of distinct α-synuclein filaments.^12, 13^

There is mounting evidence that suggests interactions between α-synuclein and lipid membranes play an important role in misfolding and aggregation of α-synuclein.^14–17^ It was previously shown that lipid vesicles consisting of anionic phospholipids (1,2-dimyristoyl-sn-glycero-3-phospho-L-serine, DMPS) promote the formation of filamentous aggregates of α-synuclein.^14^ In addition, recent high-resolution imaging of Lewy bodies (LB) extracted from PD patients revealed the co-presence of α-synuclein aggregates and lipid-rich inclusions in the LB,^18^ supporting the important role of lipids. In this study, structural features of α-synuclein filaments formed in the presence of lipid vesicles (DMPS) were investigated using cryo-electron microscopy (cryo-EM) to compare their structural characteristics to those of previous in vitro and ex vivo α-synuclein filaments.

## Results

Model membrane vesicles consisting of DMPS have previously been used to explore the effect of lipid vesicles on α-synuclein aggregation and morphology of α-synuclein aggregates mainly at pH of 6.5 and 30 °C.^14, 19–21^ In this study, additional incubation conditions were used to prepare filamentous aggregates suitable for structural characterization using cryo-EM. Small unilamellar vesicles (SUV) with an average diameter of 75 nm were prepared from DMPS via extrusion methods (Figure S1). Aggregation kinetics of wild-type (WT) and mutant (G51D) forms of α-synuclein in the presence and absence of DMPS lipid vesicles were examined at different pH conditions (pH 6.5 and 7.4) using thioflavin T (ThT) fluorescence (Figure 1). The protein aggregation was greatly accelerated in the presence of DMPS-SUV for both WT (red) and G51D (black) α-synuclein at pH 6.5 under the quiescent conditions (Figure 1a), which is consistent with previously reported kinetics monitored at pH 6.5 and 30 °C.^14,21^ In addition, WT synuclein (red) exhibits accelerated aggregation kinetics than the G51D mutant (black) at the two pHs and temperatures, presumably due to less favorable interaction between the more negatively charged G51D mutant and negatively charged DMPS lipid vesicles. The DMPS lipid vesicles induce aggregation of WT (red) and G51D mutant (black) α-synuclein at the physiological pH of 7.4 and 37 °C as well (Figure 1b).

**Figure 1.**
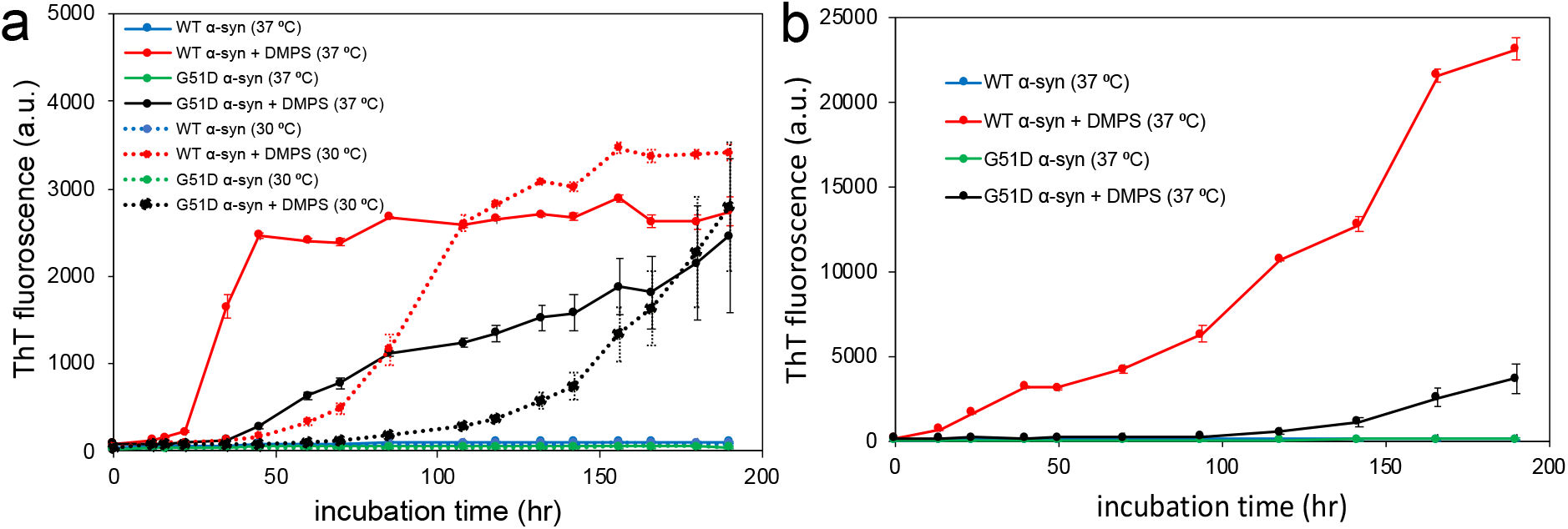
Aggregation kinetics of α-synuclein (60 μM) in the absence and presence of DMPS SUVs (100 μM) at pH 6.5 (a) and pH 7.4 (b) under quiescent conditions.

Morphological features of the aggregates were examined using transmission electron microscopy (TEM) (Figure 2). After 7 days of incubation, oligomeric species were observed for G51D α-synuclein (60 μM) in the presence of the lipid vesicles (100 μM) at pH 6.5 and 30 °C (Figure 2a). On the other hand, WT α-synuclein forms curly protofilaments at the same condition (Figure 2b). At a higher temperature of 37 °C, the flexible filamentous aggregates were observed for both G51D and WT α-synuclein at pH 6.5 (Figure 2c and 2d, respectively).

**Figure 2.**
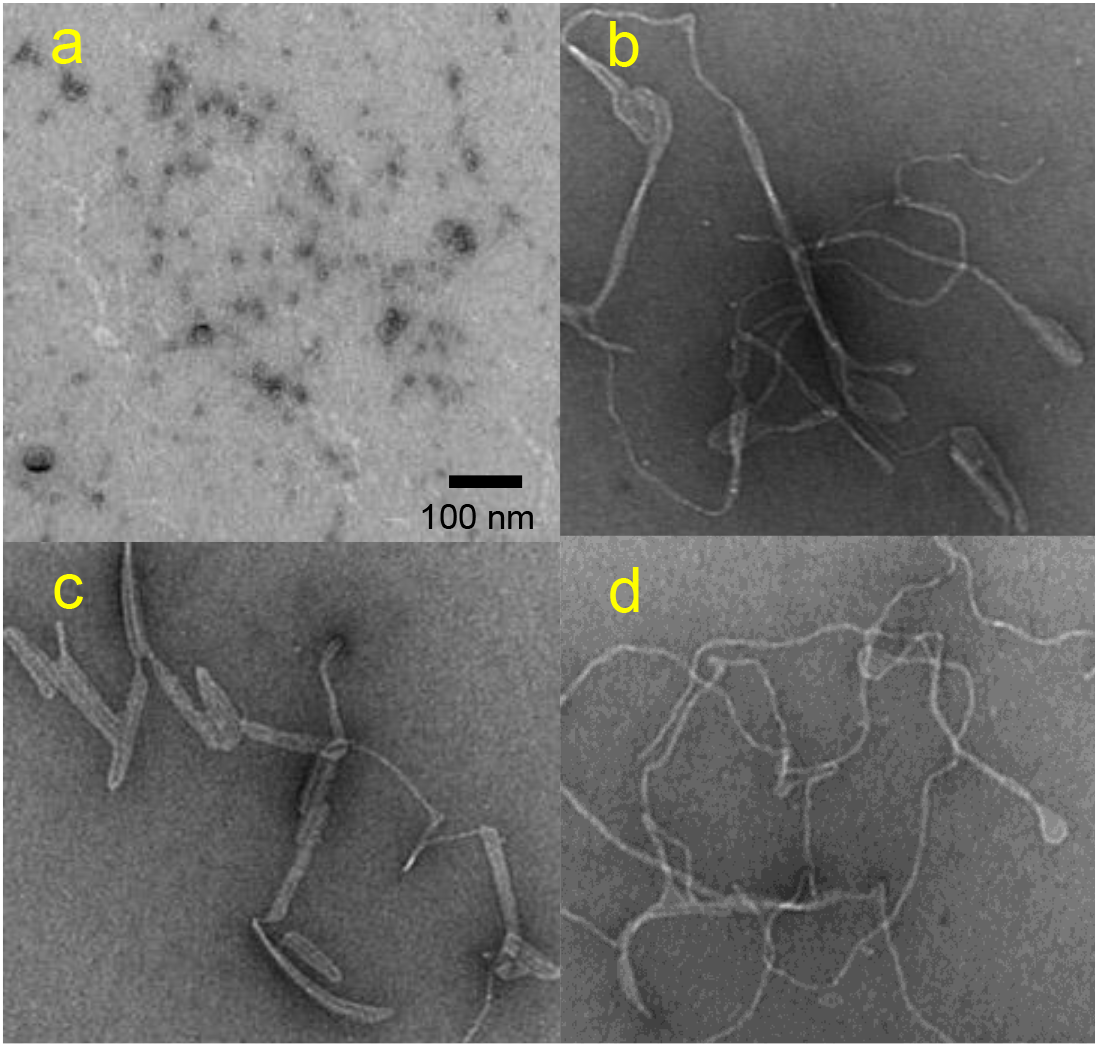
TEM images of G51D α-synuclein at pH 6.5 and 30 °C (a), WT α-synuclein at pH 6.5 and 30 °C (b), G51D α-synuclein at pH 6.5 and 37 °C (c), and WT α-synuclein at pH 6.5 and 37 °C (d). The protein samples (60 μM) were incubated for 7 days in the presence of DMPS (60 μM) lipid vesicles of an average diameter of 75 nm.

The morphology of α-synuclein filaments formed at the physiological pH (7.4) at 37 °C was also investigated by TEM (Figures S2 and S3). The mutant (G51D) filaments appear to be greatly extended at the higher pH (Figure S2). WT α-synuclein filaments also become somewhat more extended after a longer incubation of 38 days, but they still exhibit curly morphological features (Figure S3). Thus, the mutant (G51D) filaments were chosen for more detailed structural investigation using cryo-EM.

The G51D α-synuclein filaments formed at pH 7.4 and 37 °C in the presence of DMPS SUVs were frozen on a Quantifoil 1.2/1.3 300 mesh grid and images were acquired at 45,000x magnification on a 200 keV Talos Arctica electron microscope equipped with a Gatan K2 Summit direct detection camera. The cryo-EM micrographs reveal the presence of distinct thin and thick α-synuclein filaments with a diameter of 5 – 13 nm (Figure 3), similar to previously observed filaments isolated from MSA brains which had a diameter of 5 – 18 nm^22^.

**Figure 3.**
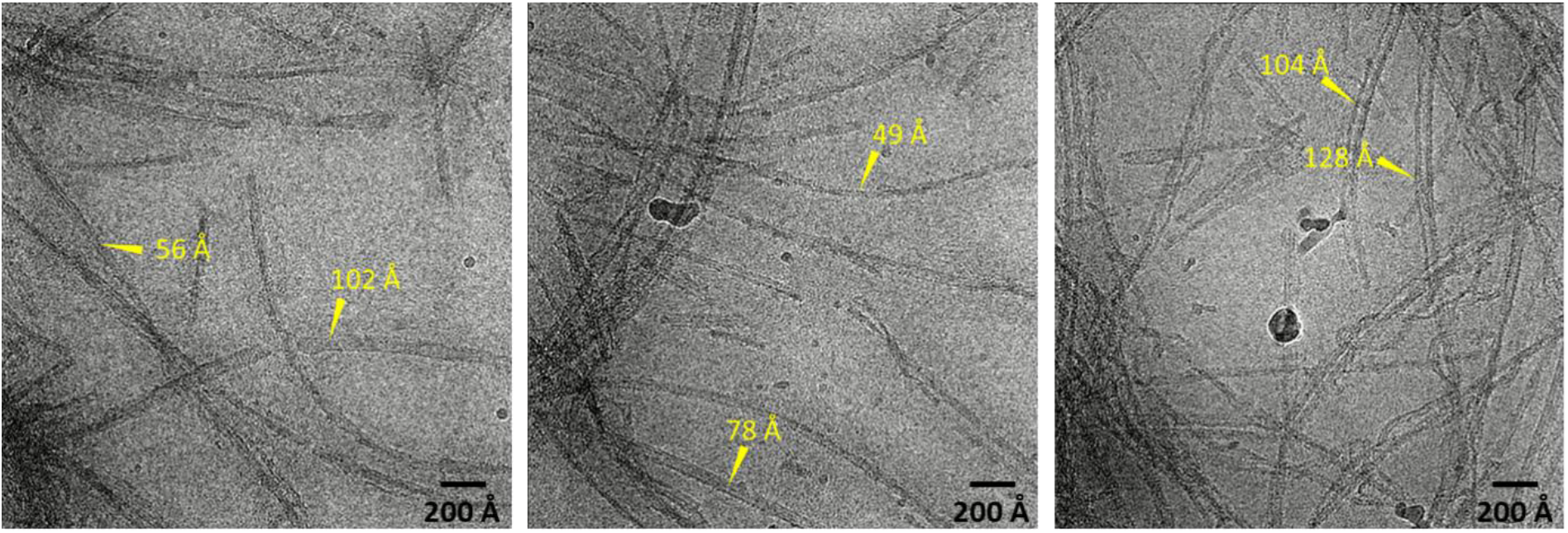
Representative cryo-EM micrographs of G51D α-synuclein (60 μM) incubated for 30 days in the presence of DMPS (100 μM) vesicles of an average diameter of 75 nm.

About 250 micrographs were analyzed using Relion reference-free two-dimensional (2D) classification, which reveals the major species of α-synuclein filaments with different thicknesses (Figure 4, Figure S4, and Table S1). The 2D class averages may originate from different orientations of single asymmetric cylindrical filament or different types of filaments. As for the asymmetric cylindrical filament, it is less likely to observe thinner images from the filaments due to the smaller surface area along that orientation. However, the thinner filaments are more frequently observed in the micrographs (Figure 3) and 2D class averages (Table S1), suggesting that α-synuclein forms at least two distinct filaments in the presence of DMPS vesicles.

**Figure 4.**
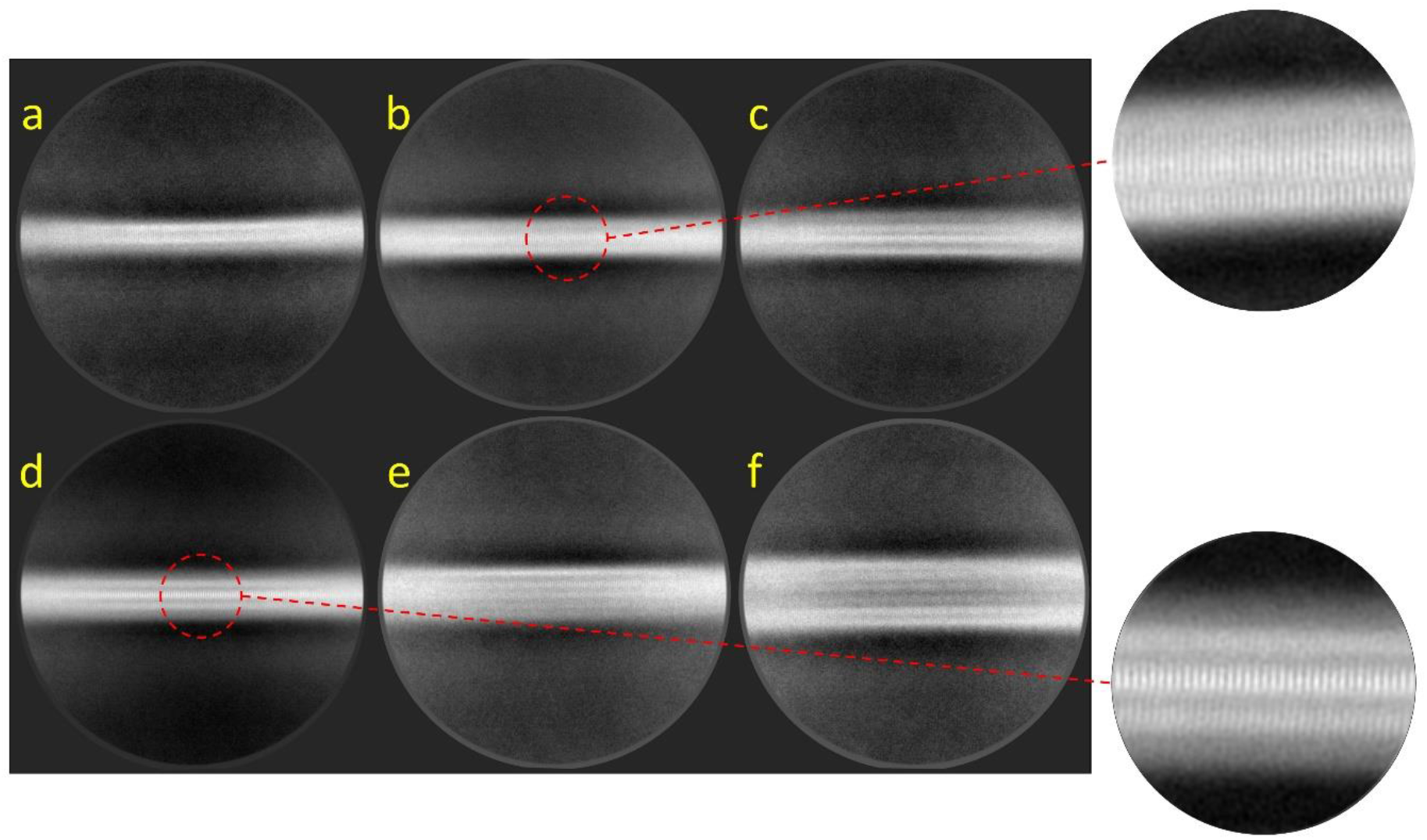
Representative reference-free 2D class averages of G51D α-synuclein filaments formed in the presence of DMPS SUVs were obtained with a box size of 56 nm.

The thin and thick filaments are stacked along the axis with a spacing of 4.7 Å, determined by the power spectrum (Figure 4 and Figure S4). It is interesting to note that the filaments do not exhibit helical twists that are observed in the previous cryo-EM structure of the in vitro filaments^2–5^ (Figure 4, Figure S4, and Table S2). These results suggest that the DMPS vesicles induce the formation of untwisted filaments, which were observed for the ex vivo filamentous aggregates extracted from DLB patients^7^.

## Discussion

Recent advances in cryo-EM and solid-state NMR methodologies enabled the structural determination of amyloid filaments at atomic resolution.^2–6^ The high-resolution structural studies revealed that α-synuclein can form diverse filamentous aggregates with different molecular conformations under different buffer conditions. Ex vivo α-synuclein filaments extracted from MSA and DLB patients’ brains were, however, shown to adopt distinct molecular conformations from those of the in vitro filaments.^7^ In particular, only untwisted filamentous aggregates were observed for the extracts from DLB patients. These results suggest the presence of co-factors involved in the formation of α-synuclein filaments in vivo.

Aggregation of α-synuclein is affected by a variety of factors such as metal ions, negatively charged lipids, other aggregation-prone proteins including tau, and DNA.^14,23–25^ Cofactors in different cellular environments may play an important role in promoting misfolding and aggregation of α-synuclein in vivo.^13^ It was also previously suggested that α-synuclein interacts with synaptic vesicle membranes in dopaminergic neurons.^26, 27^ In addition, co-localization of lipid membranes and α-synuclein aggregates in LB^18^ suggests a critical role of lipid membranes in misfolding and aggregation of α-synuclein.^28^

Model membrane vesicles consisting of DMPS have been used to explore the effect of the lipid vesicles on α-synuclein aggregation and morphology of α-synuclein aggregates.^14, 19, 20^ In this study, cryo-EM was used to investigate structural features of α-synuclein aggregates formed in the presence of DMPS vesicles. Our structural analyses revealed that α-synuclein forms untwisted filaments with different diameters of 5 – 13 nm, which are similar to those observed in brain extracts from DLB patients. However, a series of large helical fibrils with diameters of 40 – 100 nm reported in a previous structural study (100 μM of WT α-synuclein with 200 μM of DMPS at pH 6.5)^19^ are not detected in our experimental conditions (60 μM of G51D α-synuclein with 100 μM of DMPS at pH 7.4). In addition, a very recent structural study demonstrated that phospholipid vesicles consisting of POPC/POPA (1:1, lipid to protein ratio of 5:1) induced the formation of twisted WT α-synuclein filaments of 10 – 15 nm in thickness with a helical pitch of 90 – 120 nm.^29^ These results indicate that the morphology of α-synuclein filaments derived by lipid vesicles may depend on the lipid composition, protein to lipid ratio, and the mutation that may modulate α- synuclein bindings^16, 30^. Further studies are, therefore, needed to investigate the effect of those factors on the filament structure.

In summary, we report untwisted straight filaments with diameters of 5 – 13 nm formed in the presence of lipid vesicles, which have not been previously observed for in vitro α-synuclein aggregates. The untwisted straight filaments are morphologically similar to those extracted from DLB patients, suggesting that lipid vesicles may play an important role in α-synuclein aggregation in vivo. The new structures induced by the lipid vesicles also imply that interactions between α-synuclein and cofactors may lead the protein to distinct misfolding and aggregation pathways, highlighting the important role of cofactors in cellular environments.

## ASSOCIATED CONTENT

### Supporting Information

Materials and methods. TEM images of α-synuclein filaments.

The following files are available free of charge.

### AUTHOR INFORMATION

**Corresponding Author**

*.

### Author Contributions

The manuscript was written through the contributions of all authors. All authors have given approval to the final version of the manuscript.

### Funding Sources

This work was supported in part by NIH R01 NS097490 (K.H.L.), R01 AG054025 (R.K.), and R01 NS094557 (R.K.) and by Intramural Research Program of the NIH; National Institute of Environmental Health Sciences Grant ZIC ES103326 (M.J.B.).

### Notes

The authors declare no competing financial interest.

### Accession Codes

α-synuclein: UniProtKB entry P37840

## ABBREVIATIONS

DMPS: 1,2-dimyristoyl-sn-glycero-3-phospho-L-serine
TEM: transmission electron microscopy
DARR: dipolar assisted rotational resonance
cryo-EM: cryo-electron microscopy
MSA: multiple system atrophy
DLB: dementia with Lewy bodies
ThT: thioflavin T
DLS: Dynamic light scattering
CTF: contrast transfer function
POPA: 1-palmitoyl-2-oleoyl-*sn*-glycero-3-phosphate
POPC: 1-palmitoyl-2-oleoyl-*sw*-glycero-3-phosphocholine

## For Table of Contents use only

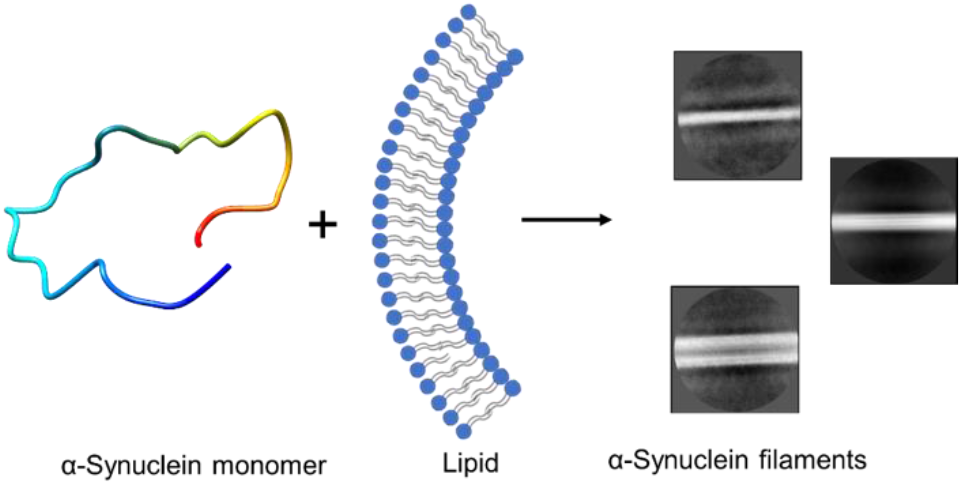

