## Supporting Information for "Untwisted α-synuclein Filaments formed in the Presence of Lipid Vesicles"

[1]Department of Chemistry, East Carolina University, Greenville, NC 27858, USA. [2]Genome Integrity and Structural Biology Laboratory, National Institute of Environmental Health Sciences, National Institutes of Health, Department of Health and Human Services, Research Triangle Park, NC, 27709, USA. [3]Institute of Molecular Biophysics, Florida State University, Tallahassee, FL 32306-4380, USA. [4]Departments of Neurology, Neuroscience and Cell Biology, University of Texas Medical Branch, Galveston, TX, 77555, USA.

### Experimental Procedures

#### α-Synuclein expression and purification

A full-length human α-synuclein plasmid (pET21a, a gift from Michael J Fox Foundation, Addgene plasmid # 51486) was transformed into BL21(DE3) *E. coli* cells, which were expressed in LB medium as described previously.[1] Briefly, the transformed *E. coli* cells were grown in LB medium containing carbenicillin (100 µg/mL) at 37 ºC. When $OD_{600}$ of the cell culture reached 0.8, α-synuclein expression was induced by adding 0.5 mM IPTG. After 12 – 16 hrs of incubation at 25 ºC, cells were harvested by centrifugation. The cell pellet was sonicated in lysis buffer (20 mM Tris, 150 mM NaCl, pH 8.0) and the soluble fraction of the cell lysate was precipitated with 50 % ammonium sulfate at 4 ºC. The protein pellet collected by centrifugation was resuspended in dialysis buffer (10 mM Tris, 2 mM EDTA, pH 8.0) and dialyzed against the same buffer overnight at 4 ºC. The protein was further purified by anion exchange chromatography (HiTrap Q HP; 20 mM Tris buffer, pH 8) followed by gel filtration (HiLoad 16/60 Superdex 75 pg) at 4 °C.

#### Preparation of lipid vesicles

1,2-dimyristoyl-sn-glycero-3-phospho-L-serine (DMPS) lipid vesicles were prepared by re-suspending dried DMPS (3 mM) in 10 mM phosphate buffer (pH 7.4) and stirring at 45 ºC for 2 hrs. The lipids were then subjected to 5 freeze/thaw cycles. Small unilamellar vesicles were prepared by extruding the lipid solution 11 times through a 0.1 µm membrane filter at 45 ºC. The average diameter of lipid vesicles was confirmed using dynamic light scattering measurements.

#### Dynamic Light Scattering (DLS):

Size distribution of the lipid vesicles was measured using DynaPro NanoStar DLS instrument. The DMPS lipid vesicles were diluted to 200 µM in 10 mM phosphate buffer (pH 7.4) and a 20 µL of lipid suspension was added to a quartz cuvette. DLS measurements were recorded with an acquisition time of 3 sec and 10 repetitions at a laser wavelength of 658 nm.

**Aggregation kinetics**

The α-synuclein aggregation kinetics were monitored with ThT fluorescence using a SpectraMax® microplate reader. α-Synuclein (60 µM) was incubated with 50 µM thioflavin T (ThT) in the presence or absence of DMPS vesicles (100 µM). A 200 µL of each sample was added in duplicates to a 96 well black clear bottom microplate and sealed airtight. Microplates were incubated at 30 ºC or 37 ºC, under the quiescent condition. ThT fluorescence emission was monitored at 482 nm with an excitation of 440 nm.

**Negative staining TEM**

The α-synuclein sample (5 µL) incubated in the presence of DMPS vesicles were added onto a carbon-coated formvar copper 300 mesh grids. After 30 sec of incubation, the excess sample was blotted off with filter paper. Grids were then stained with 5 µL of 1% uranyl acetate and incubated for 30 sec. Excess stain was blotted off with filter paper and grids were allowed to air dry. Images were acquired using Philips CM12 transmission electron microscope at an accelerating voltage of 80 kV.

**Cryo-EM sample preparation, data collection, and image processing**

A 3 µL of α-synuclein sample was applied to a glow discharged UltrAuFoil 1.2/1.3 300 mesh grid (Quantifoil) and reverse blotted for 20 sec on a Leica plunge freezing instrument with chamber conditions of 15 °C and 95 % humidity. A total number of 1050 images were collected at a nominal magnification of 45,000X on a 200 kV Talos Arctica equipped with a K2 Summit direct electron detector (Gatan). Movies were collected as a series of 60 frames at a dose rate of 6 $e^-/Å^2$/sec over 9 seconds. Beam-induced motion and drift were corrected using MotionCor2.[2] Aligned (non-dose-weighted) micrographs were used for contrast transfer function (CTF) estimation of each micrograph using GCTF.[3] The α-synuclein filaments were manually picked and extracted with different box sizes in Relion 3.1.[4] The reference-free 2D classifications were performed with 50 classes.

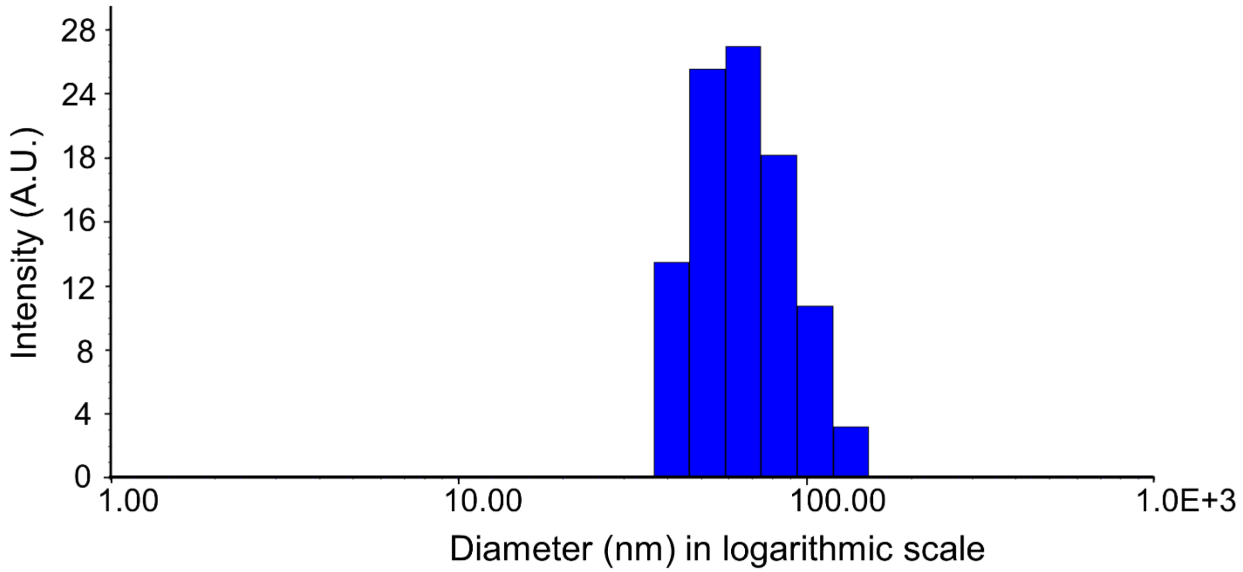


**Figure S1**. Size distribution chart of the DMPS lipid vesicles measured by DLS with a laser wavelength of 658 nm.

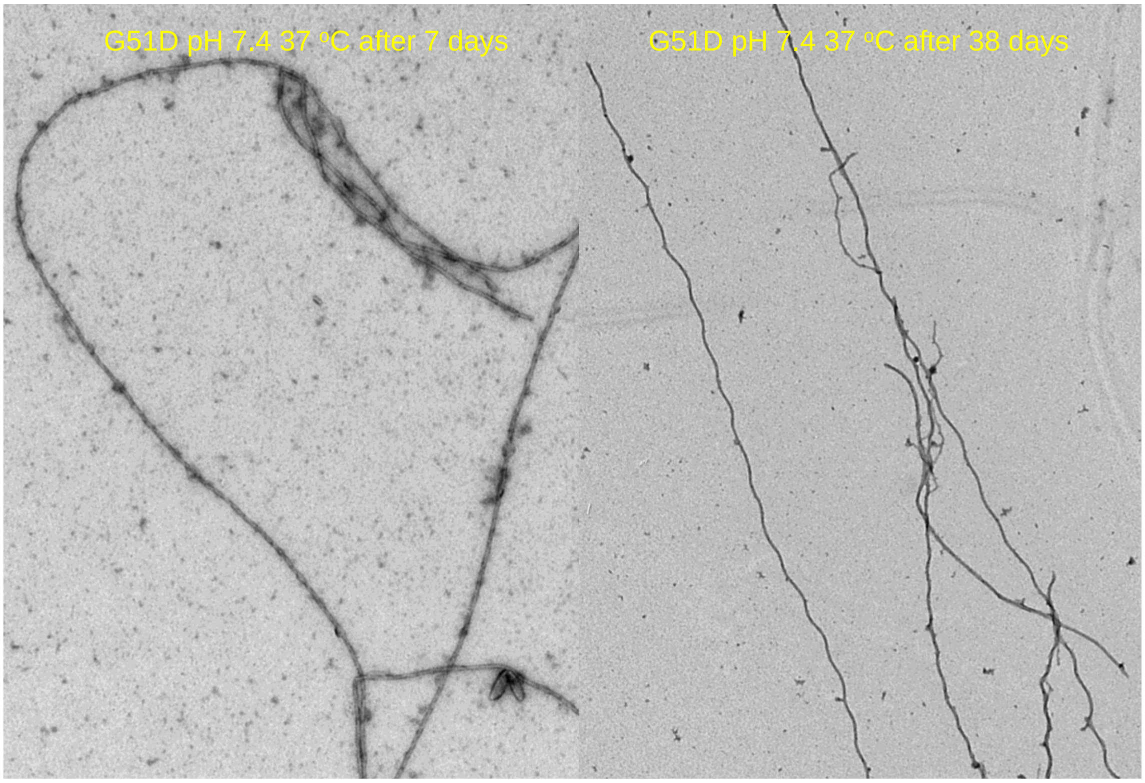


**Figure S2**. TEM images of G51D α-synuclein filaments formed in the presence of DMPS (100 μM) vesicles with a diameter of 75 nm at pH 7.4.

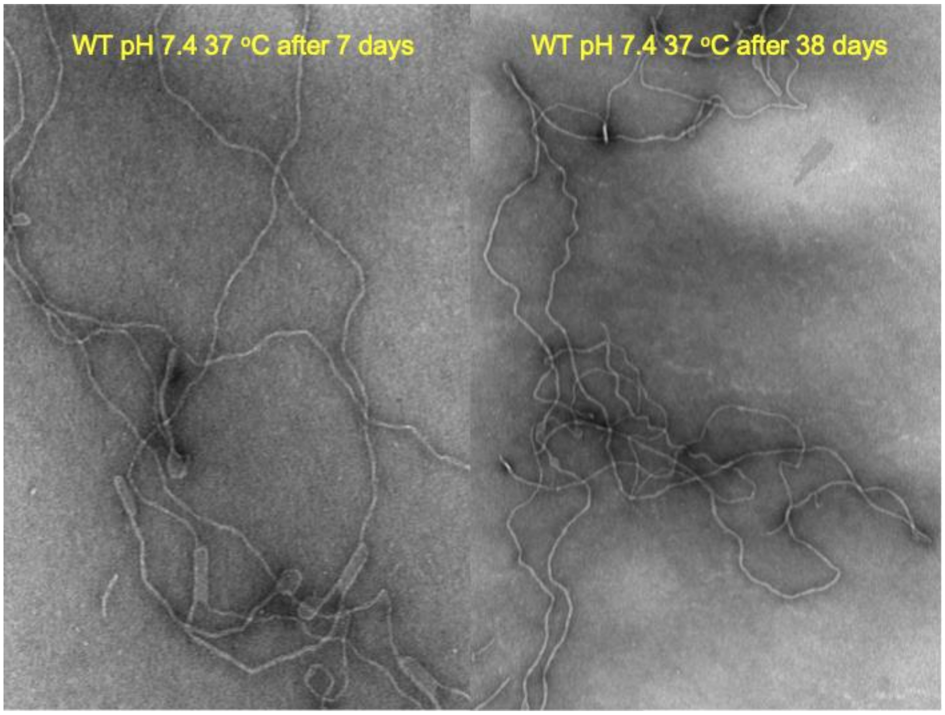


**Figure S3**. TEM images of WT α-synuclein filaments formed in the presence of DMPS (100 μM) vesicles with a diameter of 75 nm at pH 7.4.

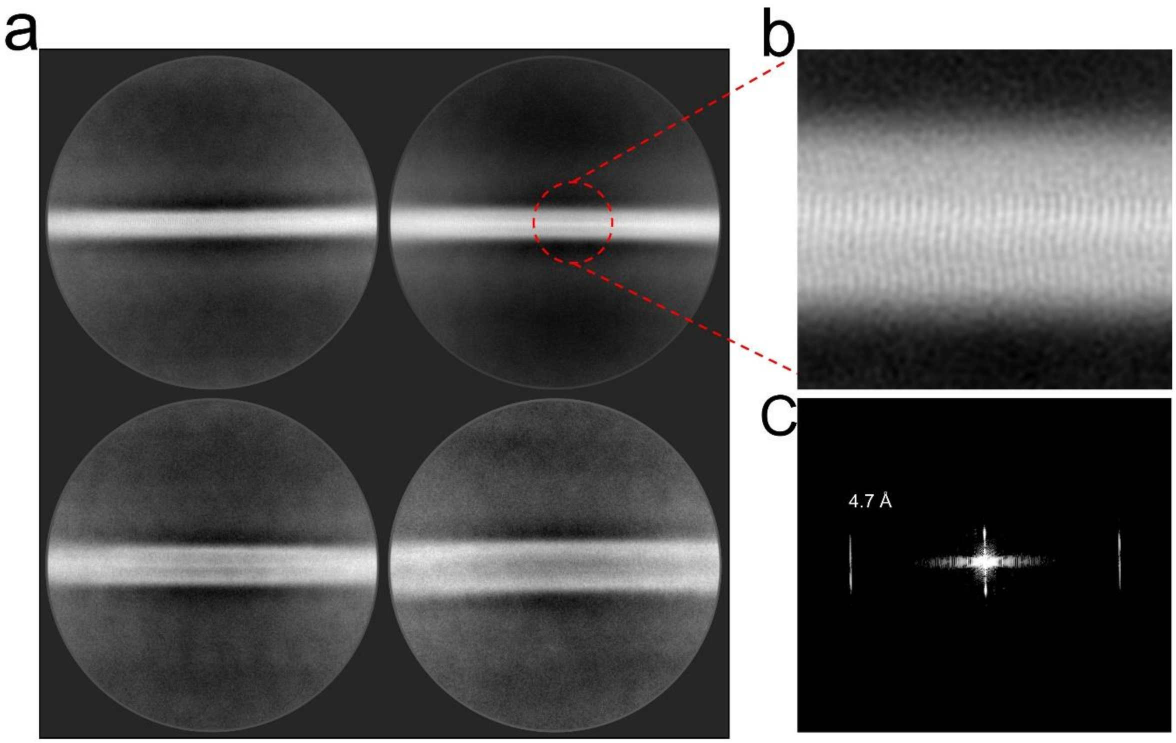


**Figure S4**. (a) Reference-free 2D class averages of α-synuclein filaments formed in the presence of DMPS SUVs obtained at a larger box size of 74 nm showing no observable twist. (b) Magnified image of a 2D class average showing the inter-strand spacing of 4.7 Å in the α-synuclein filament. (c) Representative power spectrum of the reference-free 2D class average.

| | | |
|---|---|---|
| 5.4 nm (724) | 6.7 nm (1969) | 7.6 nm (769) |
| 7.6 nm (2625) | 9.1 nm (1114) | 13.0 nm (643) |

**Table S1.** Thickness (in nm) of α-synuclein filaments for each 2D class average in Figure 4. The number of particles for each 2D class average is enclosed in parenthesis.

| PDB ID | Half-pitch (nm) |
|---|---|
| 6A6B | 120 |
| 6OSJ | 121 |
| 6CU7 | 92 |
| 6SSX | 108 |
| 6SST | 96 |
| 7L7H | 63 |
| 6XYO | 80 |
| 6XYP | 80 |
| 7NCA | 80 |
| 7NCG | 90 |

**Table S2.** Helical twists of previously reported polymorphs of α-synuclein filaments: polymorph 1a (6a6b, 6osj, 6cu7)[5,6], polymorph 2a (6ssx)[7], polymorph 2b (6sst)[7], tau-promoted polymorph (7l7h)[8], MSA type 1 (6xyo)[9], MSA type 2 (6xyp)[9], MSA seeded type 1a (7nca)[10], and type 2a (7ncg)[10].
